# Neutrophils are required during immunization with the pneumococcal conjugate vaccine for protective antibody responses and host defense against infection

**DOI:** 10.1101/2020.02.04.934380

**Authors:** Essi Y. I. Tchalla, Elizabeth A. Wohlfert, Elsa N. Bou Ghanem

## Abstract

Neutrophils can shape adaptive immunity, however their role in vaccine-induced protection against infections *in vivo* remains unclear. Here, we tested their role in the clinically relevant polysaccharide conjugate vaccine against *Streptococcus pneumoniae* (pneumococcus). We antibody depleted neutrophils during vaccination, allowed them to recover, and four weeks later challenged mice with pneumococci. We found that while isotype-treated vaccinated controls were protected against an otherwise lethal infection in naïve mice, full protection was lost upon neutrophil depletion. Compared to vaccinated controls, neutrophil-depleted mice had higher lung bacterial burdens, increased incidence of bacteremia and lower survival rates. Sera from neutrophil-depleted mice had less anti-pneumococcal IgG2c and IgG3, were less efficient at inducing opsonophagocytic killing of bacteria by neutrophils *in vitro* and worse at protecting naïve mice against pneumococcal pneumonia. In summary, neutrophils are required during vaccination for optimal host protection, which has important implications for future vaccine design against pneumococci and other pathogens.

## Introduction

*S. pneumoniae* are Gram-positive bacteria with >90 serotypes based on capsular polysaccharides [1]. These bacteria can cause pneumonia, meningitis and bacteremia [2] and remain a serious cause of mortality and morbidity worldwide, particularly in the elderly [3]. Currently, two vaccines covering common disease-causing bacterial serotypes, are available [4]. The pneumococcal polysaccharide vaccine (PPSV) consists of polysaccharides that directly cross-link B cell receptors on mature B cells leading to antibody (Ab) production independent of T cells [5]. PPSV is routinely recommended for elderly individuals >65 years old and adults with medical conditions [6]. As children <2 years old lack mature B cells, they fail to produce T-independent Abs [7]. Therefore, the pneumococcal conjugate vaccine (PCV), was introduced for use in children. PCV contains polysaccharides linked to a carrier protein that triggers a T-dependent Ab response [4]. PCV has had great efficacy in children and is currently recommended for use in immunocompromised adults and elderly individuals with underlying conditions [6]. As PCV is recommended for adults with compromised immunity including B and T cell responses [4], it is important to elucidate novel players in vaccines that could be potential targets to boost protection.

Abs against capsular polysaccharides following vaccination bind to *S. pneumoniae* and protect the host against infection [4]. The functionality of Abs is determined by their antigen affinity. Affinity to antigens is mediated via the variable regions which make up the Fab or antigen binding portions and is optimized by somatic hypermutation (SHM) [8]. The Fc or constant region of Ab, which determines their class, also shapes their function, with the different classes of Abs having distinct immune modulating activities [8]. Abs against T-independent antigens such as bacterial polysaccharides are typically produced by marginal zone B cells in the spleen [9]. In contrast, T cell-dependent Ab production occurs in germinal centers, where a specialized subset of CD4+ T-follicular helper cells (TFH) [10] induce B cells to undergo classs-witch recombination and SHM resulting in Abs with improved function [11]. PCV significantly boosts class switching to IgG as compared to PPSV [12] and further induces TFH cells which correlate with enhanced Ab function [13].

Polymorphonuclear leukocytes (PMNs) or neutrophils, play a crucial role in innate immunity to infections [14]. It is now appreciated that PMNs can also regulate adaptive immunity. PMNs can directly induce Ab production by B cells [11]. In the spleen, a subset of PMNs termed B helper neutrophils was described to produce APRIL, BAFF and IL-21 [9] that triggered Ab production by marginal-zone B cells [9, 15]. This was described for T-independent antigens including bacterial polysaccharides [9]. PMNs may also affect T cell dependent Ab responses [15], however, that is less established. PMNs are thought to both activate and suppress T cells [16]. PMNs produce a repertoire of chemokines that recruit T-cells and also produce cytokines that drive T-cell subset differentiation [16]. PMN-derived products can prime T cells to more efficiently respond to antigens [17]. PMNs also activate T cells via recruiting and activating antigen presenting cells [18] or acting as antigen presenting cells themselves [16, 19-23]. In contrast, PMNs produce compounds that inhibit T cell activation [24] including ROS [25], arginase-1 [26] and serine proteases [27]. Therefore, it is unclear whether upon *in vivo* vaccination, if PMNs would suppress or induce T-dependent Ab responses. Further, although there have been elegant studies characterizing mechanisms of PMN interactions with B and T cells, most of the work has been done either *in vitro* or *in vivo* using model antigens [11, 18, 24]. Thus, studies examining the role of PMNs in clinically relevant vaccinations and how that shapes protection against *in vivo* infections are needed.

PMNs are required to control bacterial numbers following *S. pneumoniae* infection [28, 29]. PMNs also play a role in Ab responses against pneumococci. When compared to healthy controls, patients with neutropenic disorders had lower levels of Abs to some pneumococcal polysaccharides [9]. In mice, splenic PMNs localized with marginal zone B cells and were required for production of T-independent Abs during pneumococcal infection [15, 30]. However, whether PMNs shape responses to the pneumococcal conjugate vaccine and if they impair or promote Ab production remains unexplored. Here we tested the role of PMNs in response to PCV and found they were required at the time of vaccination for optimal Ab responses as well as host protection against pneumococcal infection. This study highlights the link between PMNs and Ab responses in the context of a clinically relevant immunization, which has far-reaching implications for vaccine design against *S. pneumoniae* and other pathogens.

## Methods

### Mice

Female C57BL/6 mice (6-8 weeks) were purchased from Jackson Laboratories (Bar Harbor, ME) and used in all experiments. Mice were housed in a specific-pathogen free facility at the University at Buffalo and all experiments were conducted in accordance with Institutional Animal Care and Use Committee (IACUC) guidelines.

### Bacteria

Wild type (WT) *S. pneumoniae* TIGR4 and capsule-deletion mutant (Δ*cps*) *S. pneumoniae* were kind gifts from Andrew Camilli. All bacteria were grown to mid-exponential phase in Todd-Hewitt broth (BD Biosciences) supplemented with Oxyrase (Oxyrase) and 0.5% yeast extract at 37°C in 5% CO_2_. Aliquots were frozen at −80°C in growth media with 20% (v/v) glycerol. Prior to use, aliquots were thawed on ice, washed and diluted in PBS to the desired concentrations. Titers were confirmed by plating on Tryptic Soy Agar plates supplemented with 5% sheep blood agar (Northeast Laboratory Services).

### Immunization

Mice were immunized via intramuscular (*i.m*.) injection of 50μl of the pneumococcal conjugate vaccine Prevnar-13® (Wyeth pharmaceuticals) into the caudal thigh muscle. Sera was collected from all mice prior to immunization, as well as two and four weeks post immunization and saved at −80°C for subsequent assays.

### Neutrophil Depletion

Mice were treated intra-peritoneally (i.p.) with 50 μg of the Ly6G-depleting antibody IA8 or isotype IgG control (BioXCell) following the timeline in Fig 1A.

**Figure 1.**
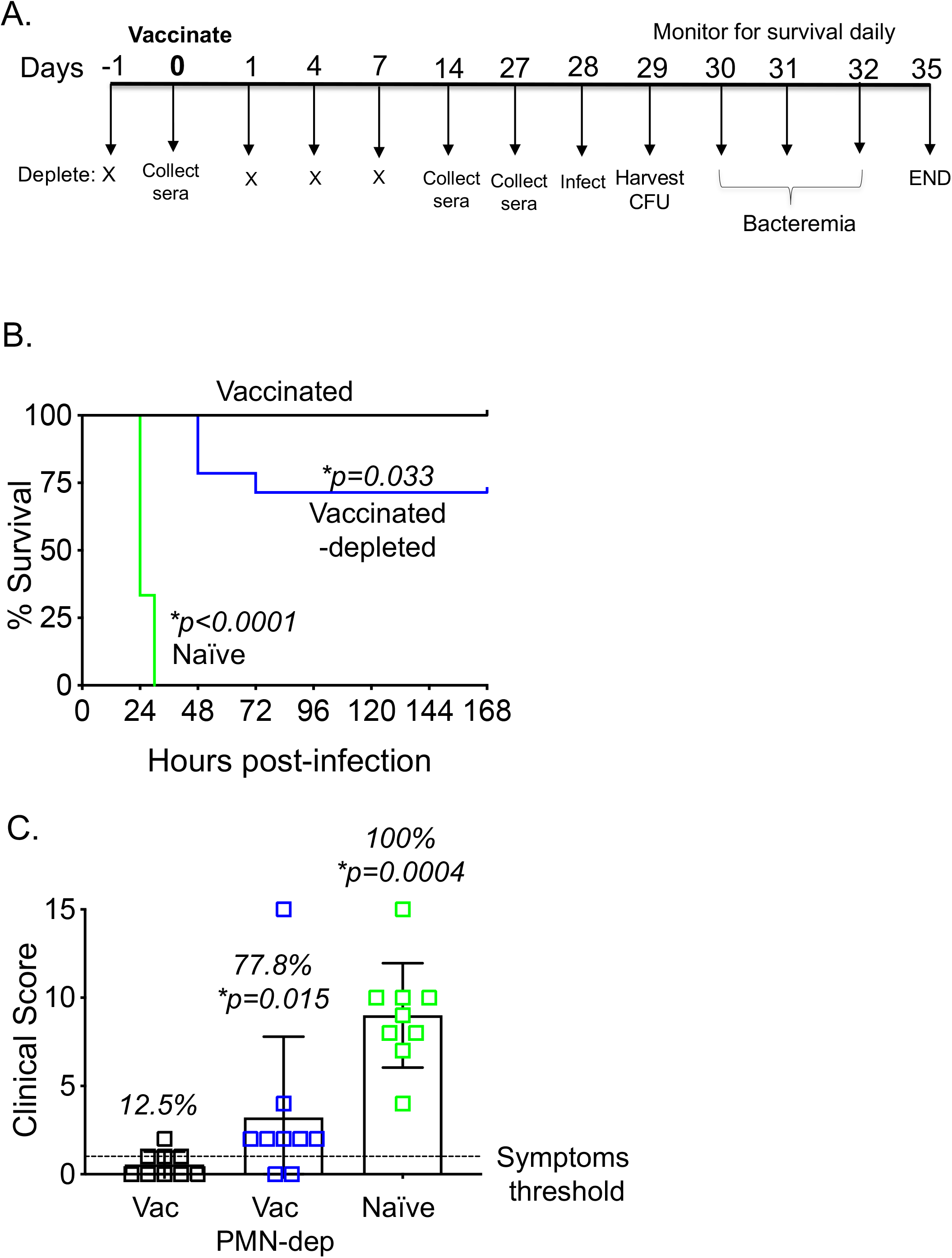
PMNs are required at the time of vaccination for PCV-mediated protection against *S. pneumoniae* infection. C57BL/6 female mice were treated i.p. with PMN depleting antibodies (IA8) or isotype control at days −1, +1, +4 and +7 with respect to vaccination following the timeline outlined in panel A. Mice were mock treated (naïve) or administered 50μl of Prevnar-13 via intramuscular injections to the hind legs (vaccinated). Four weeks following vaccination mice were challenged i.t. with 1×10^7^ CFU *S. pneumoniae* TIGR4 and monitored for survival over time (B) and clinical signs of disease (C). (B) Data were pooled from 14 mice/group from three separate experiments and * denotes significance calculated by the log-Rank (Mantel-Cox) test. (C) Data were pooled from three separate experiments with each square representing an individual mouse. The dashed line indicates the symptomatic score threshold (above one). Fractions indicate the percent of mice that had a score above 1 and * denotes significant differences from vaccinated controls by Fisher’s exact test.

### Adoptive Transfer of Sera

Five weeks following immunization, vaccinated, vaccinated PMN-depleted and naïve mice were euthanized and blood harvested via cardiac puncture. Sera was obtained from the blood, pooled for each group and transferred i.p (250μl) into naïve recipients. Recipients were then infected one hour later [31].

### Animal Infections

Mice were intra-tracheally (i.t.) challenged with 107 colony-forming units (CFU) of WT *S. pneumoniae* as previously described [31]. Following infection, one set of mice were monitored daily over one week for bacteremia as well as clinical signs of disease including weight loss, activity level, posture and breathing and blindly scored from 0 (healthy) to 21 (severely sick). Twenty-four hours post infection, another set of mice were euthanized and lung and blood were assessed for CFU.

### Antibody ELISA

Sera Ab levels were measured by ELISA as previously described [31]. Nunc maxisorp® plates were coated overnight at 4°C with type 4 Pneumococcal Polysaccharide (ATCC®) at 2ug/well. Plates were washed and blocked for 2 hours. The sera were preabsorbed with a pneumococcal cell wall polysaccharide mixture (CWP-multi from Cederlane) to neutralize non-capsular Abs and then added to the plate. After a 3h incubation and washing, pneumococcal-specific Abs were detected using HRP-conjugated goat anti-mouse IgM (Invitrogen), IgG (Millipore Sigma), IgG1, IgG2b, IgG2c or, IgG3 (Southern Biotech) followed by TMB substrate (Thermo Scientific^™^) and readings at OD_650_ using a BioTek^®^ reader. Kinetic ELISAs were performed with readings every minute for 10 minutes. Ab units were calculated as percentages of a control hyperimmune serum included in each ELISA. Hyperimmune sera was pooled from mice that were intra-nasally inoculated with *S. pneumoniae* TIGR4 over four weeks as previously described [31], immunized with PCV at week 4 and injected i.p. with heat-killed bacteria at week 5.

### Myeloperoxidase (MPO) ELISA

MPO levels were measured in lung homogenates using the Mouse Myeloperoxidase ELISA kit from InvitrogenTM as per manufacturer’s instructions.

### Isolation of PMNs

PMNs were isolated from the bone-marrow using density centrifugation with Histopaque 1119 and Histopaque 1077 (Sigma) as previously described [28]. PMNs were resuspended in Hanks’ Balanced Salt Solution (HBSS without Ca2+ and Mg2+) supplemented with 0.1% gelatin and used in subsequent experiments. Purity was confirmed by flowcytometry and the isolated cells were 85-90% Ly6G+.

### Opsonophagocytic Killing Assay (OPH)

The ability of PMNs to kill pneumococci was assessed as previously described [28]. Briefly, 1×10^5^ PMNs were infected with 1×10^3^ bacteria pre-opsonized with 3% mouse sera in 100 μl reaction volumes of HBSS/0.1% gelatin (with Ca2+ and Mg2+) and rotated at 37°C for 40 minutes. The reactions were stopped on ice and plated for CFU. The percent of bacteria killed was calculated using no PMN controls.

### Flow Cytometry

One day following the last PMN depletion, mice were euthanized and blood, vaccine draining popliteal lymph nodes and spleen were harvested. Single-cell suspensions of splenocytes and lymph nodes were prepared by mashing the organs through sterile mesh screens using the plunger of a 3-ml syringe. Red blood cells were lysed with a hypotonic buffer and the cells surface stained for Ly6G (IA8 or RB6, Biolegend), CD11b (M1/70, Invitrogen,), CD11c (N418, BD Bioscience), F4/80 (BM8, BD Bioscience) and Ly6C (AL-21. BD Bioscience) in the presence of Fc-block (BD Bioscience). Live cells were identified using a dead cell stain kit (Life Technologies). Fluorescence intensities were measured on a BD Fortessa and at least 20,000 events were analyzed using FlowJo.

### Statistical Analysis

All statistical analysis was done using Graphad Prism version 8. Significant differences were determined by Fisher’s exact test, One-way ANOVA followed by Dunnet’s test or Student’s t-test as appropriate. Survival analyses were performed using the log-rank (Mantel-Cox) test. *p* values less than 0.05 were considered significant.

## Results

### PMNs are required at the time of immunization with PCV for host protection against pneumococcal infection

To test if PMNs were required for protection at the time of vaccination with PCV, we used the anti-Ly6G Ab IA8 to deplete PMNs or isotype controls one day prior to and every two days throughout the first week following vaccination (timeline- Fig 1A). One day after the final treatment, we verified depletion in the blood and found ~99% reduction in the number of circulating PMNs (Fig S1A). As splenic PMNs have a role in Ab production [9, 15], we also examined PMN numbers in the spleen. We found that upon vaccination there was a ~2-fold increase in splenic PMNs (Fig S1B) and that treatment with depleting Abs resulted in ~98% depletion of those cells (Fig S1B). We also verified that Ab treatment was specific to PMNs and did not result in any changes in the number of circulating and splenic monocytes, dendritic cells or macrophages (Fig S2).

Four weeks following vaccination, we challenged mice i.t. with *S. pneumoniae* TIGR4 strain. Invasive *S. pneumoniae* infection results in pneumonia primarily, but up to 30% of patients with pneumococcal pneumonia also develop bacteremia and have a worse prognosis [32]. As *S. pneumoniae* strains can differ considerably [33], we chose the well-characterized serotype 4 isolate TIGR4, originally isolated from a bacteremic patient, as a model of a highly invasive infection modeling pneumonia that results in bacteremia [34] and that is covered by PCV. We then monitored the disease course and as expected found that while all naïve mice rapidly succumbed to infection, 100% of vaccinated mice survived (Fig 1B). However, unlike vaccinated controls, full protection was lost in PMN-depleted mice (Fig 1). The majority of PMN-depleted mice displayed severe clinical signs of disease where 77.8% got sick as compared to only 12.5% of vaccinated controls (Fig 1C). PMN-depleted mice had between 10-100-fold higher pulmonary bacterial numbers (Fig 2A) and systemic spread (Fig 2B) into the circulation, culminating in significantly reduced survival (Fig 1B) as compared to vaccinated controls. This reduced protection in PMN-depleted mice was not due to the continued absence of PMNs at the time of challenge, as we verified that there was no difference in PMN presence in the lungs as measured by MPO levels following infection (Fig S1C). Rather, our findings suggest that PMNs are required at the time of vaccination with PCV for full protection against subsequent pneumococcal infection.

**Figure 2.**
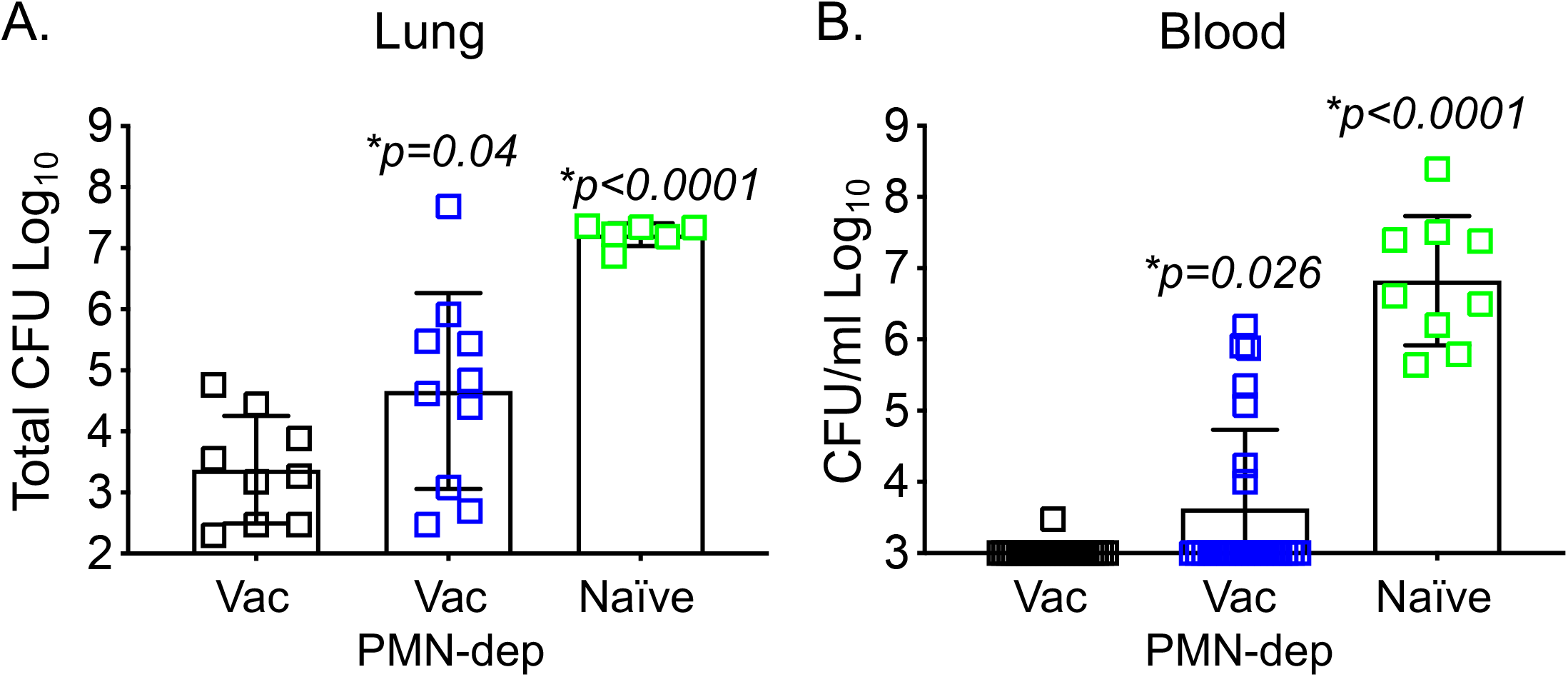
PMNs are required at the time of PCV vaccination for subsequent control of *S. pneumoniae* burden upon pulmonary challenge. Naïve (green), Prevnar-13 immunized (black) and PMN depleted Prevnar-13 immunized mice (blue) were challenged i.t. with 1×10^7^ CFU *S. pneumoniae* TIGR4 four weeks following vaccination following the timeline in Fig1A. Bacterial burden in the lungs (A) and blood (B) were also enumerated 24 hours post infection. Data were pooled from three separate experiments with each square representing an individual mouse. * denotes significant differences from vaccinated controls by One-way ANOVA followed by Dunnet’s test.

### PMNs are required for optimal antibody isotype switching in response to PCV immunization

Next, we wanted to explore the mechanisms by which PMNs contributed to vaccine-induced protection. We first examined Ab production and as expected observed isotype switching to IgG by week 4 post vaccination in our control group (Fig 3). We found that PMN depletion did not alter IgM or total IgG levels against capsular polysaccharide type 4 (Fig 3A and B) or heat-killed *S. pneumoniae* (not shown). However, when we examined IgG subtypes, we found that depletion of PMNs during vaccination resulted in slightly reduced levels of IgG2b (Fig 4A) and significantly lower levels of IgG2c (Fig 4B) and IgG3 (Fig 4C) at week 4 post vaccination. Interestingly, IgG1 levels (Fig 4D) were slightly, but not significantly elevated in PMN depleted group as compared to vaccinated controls. These data suggest that PMNs play a role in class switching to certain IgG subtypes.

**Figure 3.**
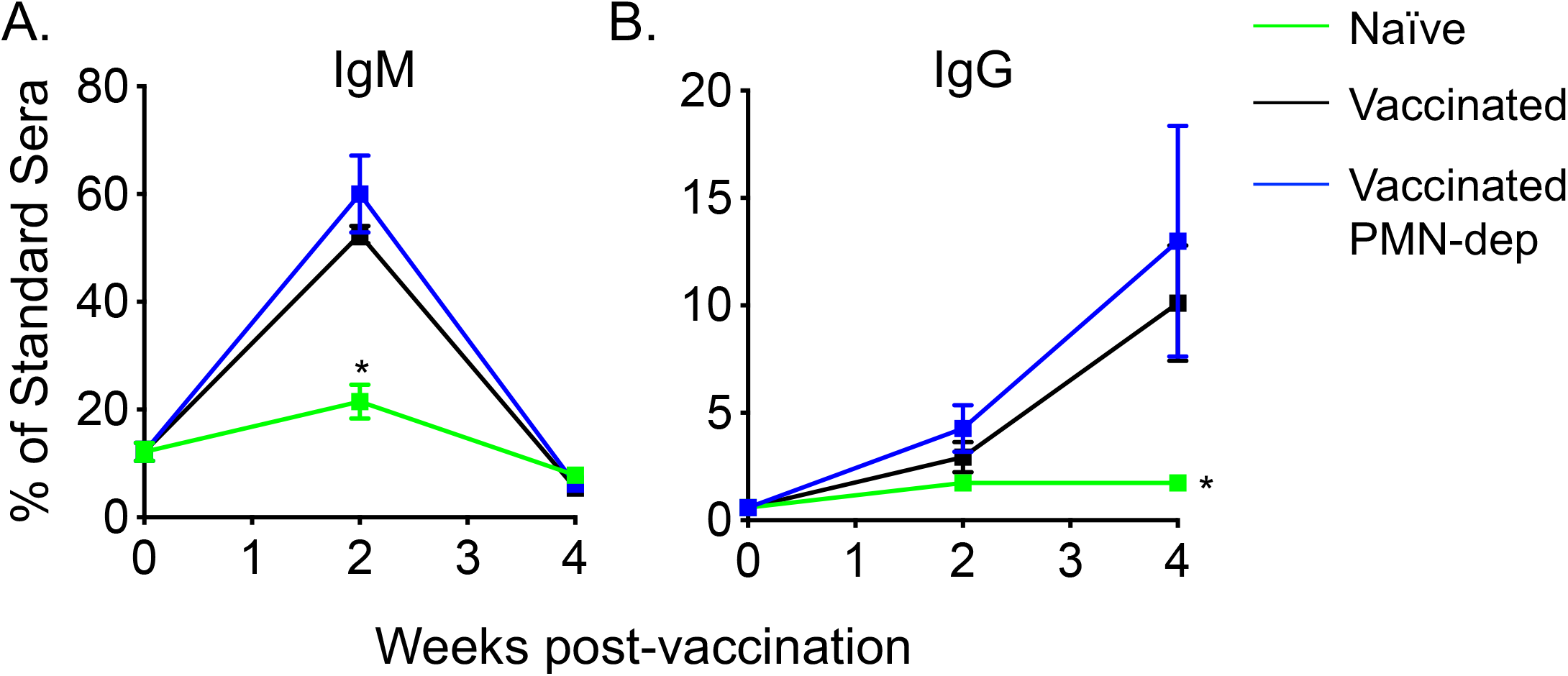
Total levels of anti-pneumococcal IgG and IgM remain unchanged in PMN-depleted PCV immunized mice. Sera were collected from naïve (green lines), Prevnar-13 immunized (black lines) and PMN depleted Prevnar-13 immunized mice (blue lines) over time as indicated in Fig 1A. Circulating levels of IgM (A) and total IgG (B) against purified polysaccharide serotype 4 were then measured by ELISA. Antibody units were calculated based on a hyperimmune standard (see Materials and Methods) included in each ELISA plate. *p* values were determined by student t-test. Asterisks (*p*<0.05) indicate significant differences with respect to vaccinated mice. Data were pooled from two separate experiments with n=6 mice per group and presented as means +/-SD.

**Figure 4.**
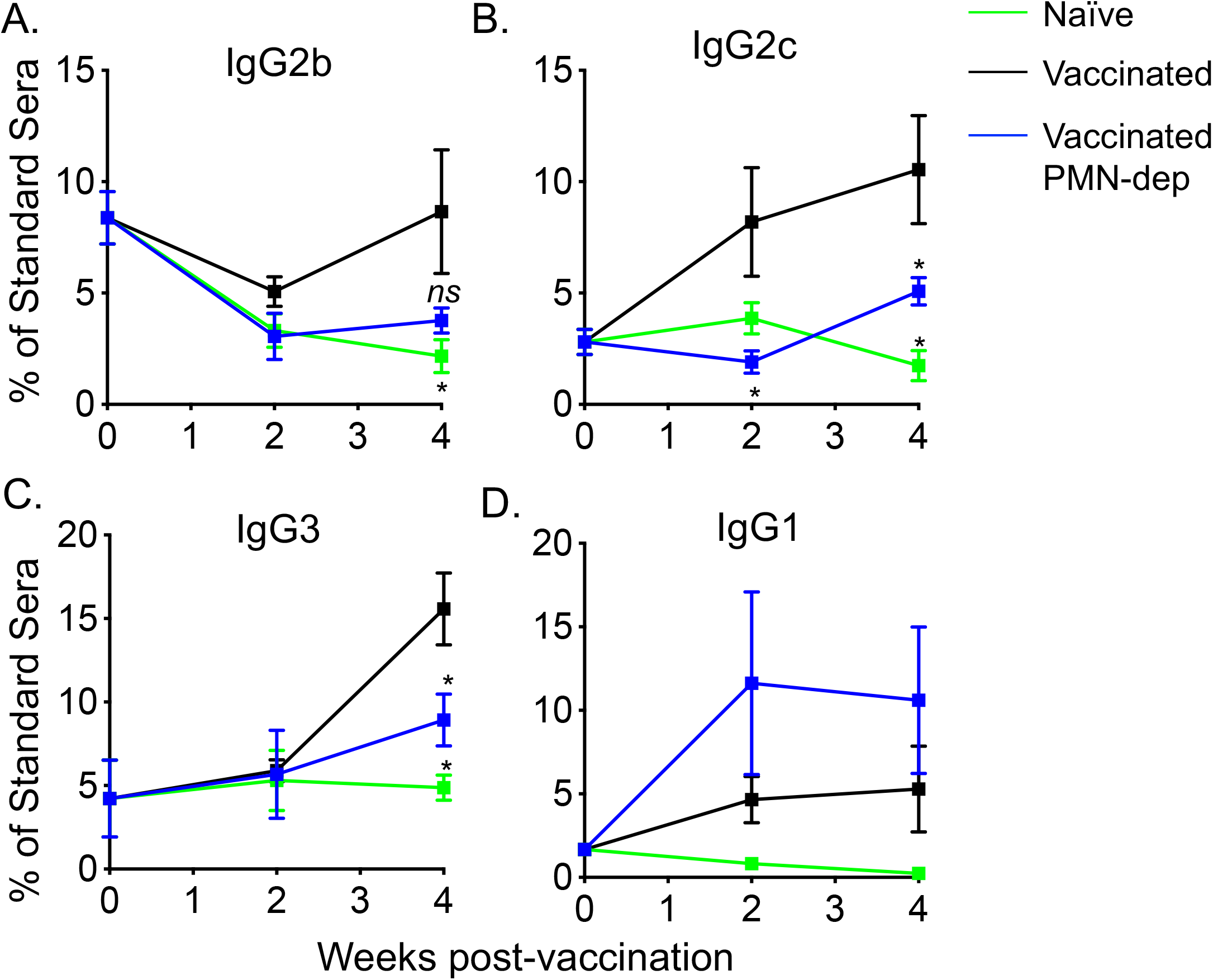
PMNs contribute to IgG2 and IgG3 production following PCV immunization. Sera were collected from naïve (green lines), Prevnar-13 immunized (black lines) and PMN depleted Prevnar-13 immunized mice (blue lines) following the timeline presented in Fig 1A. (A-D) The levels of the indicated antibodies against purified polysaccharide serotype 4 were then measured in the sera by ELISA. Antibody units were calculated based on a hyperimmune standard. *p* values were determined by student t-test. Asterisks (*p*<0.05) indicate significant differences with respect to vaccinated mice. Pooled data from two separate experiments with n=6 mice per group are presented as means +/-SD.

### PMNs are required for optimal antibody function following PCV immunization

Apart from Ab levels, Ab function is key for vaccine-efficacy [35]. We next explored if PMNs affected Ab affinity to bacterial surfaces. We tested the ability of IgG in the sera of the different mouse groups to bind the surface of *S. pneumoniae* by flow cytometry. Very little IgG bound to bacteria upon incubation with naïve sera. However, we observed a 30-fold increase in the amount of IgG bound to bacteria when sera from vaccinated mice were used (Fig 5A). As expected, in immune sera, the bound IgG was specific to capsular polysaccharides as very little IgG bound to acapsular bacteria (*Δcps S. pneumoniae*). Interestingly, we observed a significant decrease in the amount of IgG bound to *S. pneumoniae* opsonized with sera from PMN-depleted mice as compared sera from vaccinated controls (Fig 5A).

**Figure 5.**
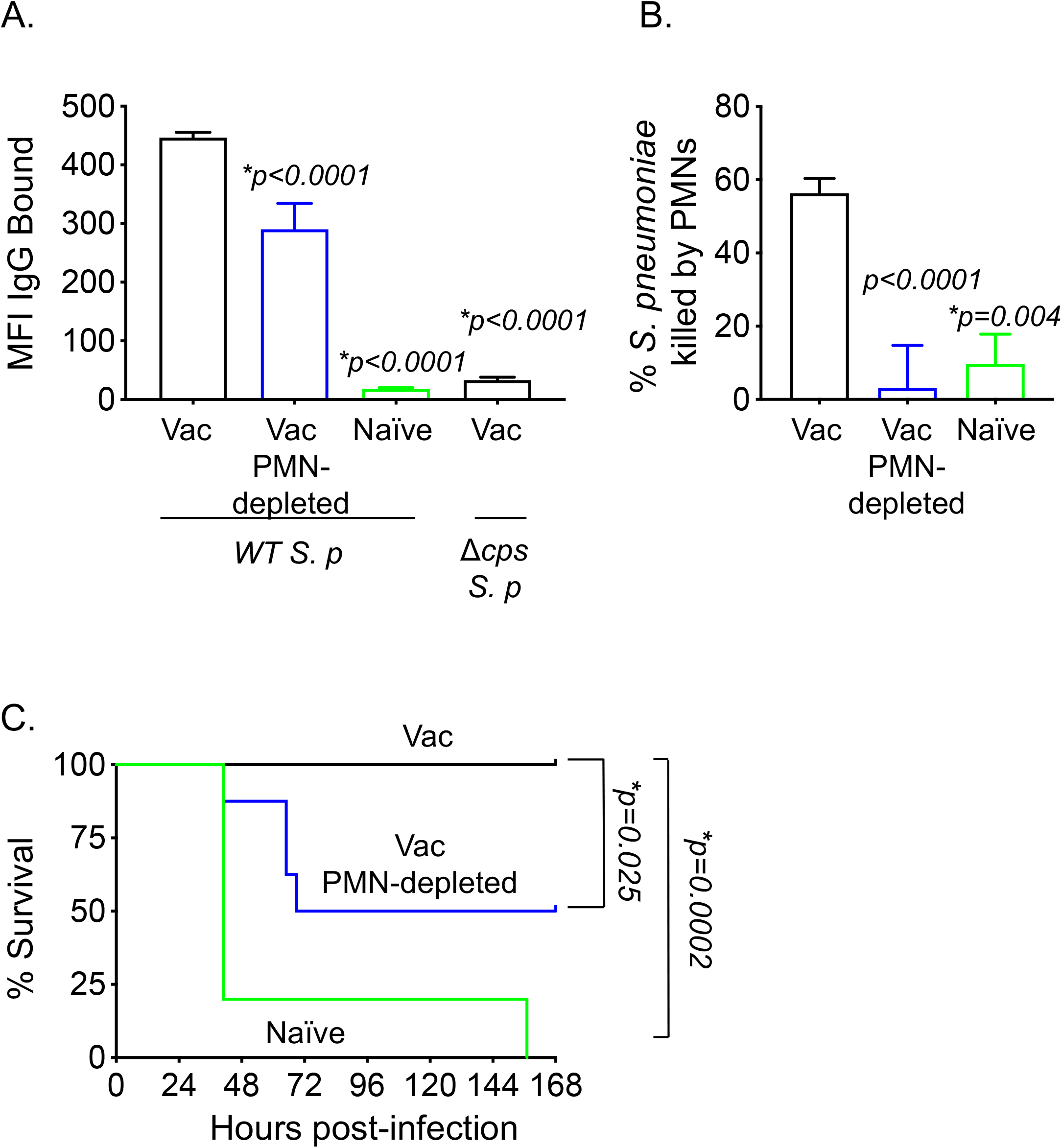
PMNs are required for optimal antibody function following PCV. (A-C) Sera were collected from naïve, Prevnar-13 immunized and PMN depleted immunized mice four weeks post vaccination following the timeline indicated in Fig 1A. (A) Wild type (*WT*) or a capsule deletion mutant (*Δcps*) *S. pneumoniae* were incubated with the indicated sera for 30 minutes, washed and stained with fluorescently-labeled anti-mouse IgG. The amount (mean fluorescent intensity or MFI) of bound Abs was determined by flow cytometry. Representative data from one of three separate experiments (n=3 biological replicates) are shown where each condition was tested in triplicate (n=3 technical replicates) per experiment. (B) The ability of PMNs isolated from naïve mice to kill pneumococci pre-opsonized with the indicated sera was determined. Percent bacterial killing was determined with respect to a no PMN control. Data shown are pooled from three separate experiments (n=3 biological replicates) where each condition was tested in triplicate (n=3 technical replicates) per experiment. (A-B) Bar graph represent means+/-SD and asterisks indicate significant differences from vaccinated controls as calculated by One-way ANOVA followed by Dunnet’s test. (C) Naïve C57BL/6 female mice were injected i.p with 200μl of pooled serum from the indicated mice then challenged i.t. 1 hour later with 5×10^5^ CFU *S. pneumoniae* TIGR4. Survival was assessed over time. *, denotes significance by the log-Rank (Mantel-Cox) test. Data were pooled from 8 mice/group from two separate experiments.

We next compared the opsonic capacity of Abs by comparing the ability of sera to induce opsonophagocytic (OPH) killing of *S. pneumoniae* by primary PMNs isolated from naive mice. We found sera from vaccinated controls significantly boosted bacterial killing by PMNs where 60% of the bacterial input were killed by PMNs in the presence of immune sera as compared to ~10% with naïve sera (Fig 5B). Strikingly, sera from PMN-depleted mice failed to induce opsonophagocytic killing of *S. pneumoniae* by PMNs where only 3% of the bacterial input was killed (Fig 5B). Abs can also activate the complement pathway and directly kill bacteria [8] but we detected no differences in the ability of sera alone from any of the mouse groups to kill pneumococci (Fig S3).

Given the difference in the *in vitro* function we observed, we finally tested the protective activity of Abs generated upon vaccination in the absence of PMNs. Naive young mice were injected i.p. with five-week sera from either vaccinated controls, naïve mock-immunized mice or our PMN depleted vaccinated group. Mice were then challenged i.t. with *S. pneumoniae* TIGR4 one hour following sera transfer. We found that while all of the mice receiving naïve sera succumbed to infection, all of the mice receiving sera from vaccinated controls survived the challenge (Fig 5C). In contrast, only half of the mice receiving sera from the PMN depleted vaccinated group survived (Fig 5C). These data indicate that Abs produced during vaccination in the absence of PMNs are not sufficient to provide protection against subsequent pneumococcal infection.

## Discussion

Traditionally, PMNs are viewed as effectors of vaccine responses where vaccination triggers Abs that bind to pathogens and promote their clearance via enhancing uptake and killing by PMNs [36]. However, the extent to which PMNs contribute to vaccine mediated protection against infections *in vivo* has not been fully elucidated. In this study, we explored the role of PMNs in immunization with the pneumococcal conjugate vaccine. We found that PMNs were needed for production of functional Abs following vaccination. Importantly, PMNs were required at the time of immunization for full protection against subsequent invasive pneumococcal infection. Our findings highlight the *in vivo* role of PMNs as inducers of protective vaccine responses against *S. pneumoniae* infections.

The mechanisms by which PMNs mediate Ab production in response to PCV is unclear. In adults, polysaccharides can directly cross-link B cell receptors and elicit Ab production independent of T cells [5]. PCV converts this T-independent response to one that involves T cells as it consists of polysaccharides linked to the carrier protein CRM197 [5]. This generates T cells specific to the carrier protein [37, 38]. When B cells recognize polysaccharides, they are thought to bind and internalize the polysaccharide along with its protein carrier and then display peptides derived from the carrier on MHC-II. This allows these polysaccharide-specific B cells to interact with carrier-peptide specific T cells which in turn help the B cells produce anti-polysaccharide Abs [39]. Therefore, in the context of PCV, PMNs could either be working on B cells, T cells or both. In humans, a subset of splenic PMNs directly induce Ab production by marginal zone B cells in response to T-independent antigens including bacterial polysaccharides [9]. In mice, splenic B helper PMNs were found to produce pantrexin3, which was important for IgM production following the immunization with the unconjugated pneumococcal polysaccharide vaccine [15]. Pantrexin3 was also important for T-cell independent IgM and IgG production against polysaccharides following intravenous infection with *S. pneumoniae* [15]. The role of PMNs in T-dependent responses is less clear. In mice, PMNs impaired IgA but not IgG or IgM production in response to vaccination with the adjuvant *Bacillus anthracis* edema toxin [40]. Mouse PMNs were found to directly present ovalbumin peptides to CD4+ T cells triggering T cell cytokine production and proliferation [20]. However, IgG2 responses to ovalbumin were not impaired in pantrexin3-/-mice [15]. In contrast, Abs against influenza PR8 and the T-dependent antigen TNP-Ficoll required pantrexin3 production by murine PMNs [15]. Similarly, human PMNs were able to present influenza hemagglutinin to CD4+ T cells [22]. In rhesus macaques, PMNs presented HIV-envelope glycoproteins to CD4+ T cells [22] and induced Ab production against SIV when co-cultured with B cells [41]. Here, in the context of immunization with PCV, PMNs are clearly required for optimal Ab responses, however whether they are acting on T cells remains to be determined.

A key finding here is that PMNs are required for the production of functional Abs. The functionality of Abs is determined by the affinity and avidity to their antigen [8]. Here, although we detected similar levels of IgM and IgG in sera from PMN-depleted and isotype-treated vaccinated mice, the ability of IgG to bind pneumococci significantly decreased when they were generated in the absence of PMNs. This suggests that Ab affinity is increased in the presence of PMNs. Ab affinity is improved by SHM of the Fab variable regions and typically occurs in germinal centers and requires T cells [42], although human splenic B helper PMNs may contribute to SHM in marginal zone B cells [9].

Ab function is also influenced by their subclass which is determined by their Fc region [8]. The Fc portion of Abs shape effector function since they determine binding to Fc receptors on PMNs as well as the ability to activate complement [8]. Here, PMNs were required for class switching to IgG2c and IgG3 but not IgG1 subtypes. The IgG subtype produced in response to PCV varies in humans based on age, with IgG2 being the predominant response in adults [12, 43]. We found that PMNs were crucial for the ability of Abs to elicit opsonophagocytic killing of bacteria by primary immune cells. This is in line with data from humans where although the opsonophagocytic activity of IgG subtypes vary based on bacterial serotype, IgG2 was reported to have the highest activity while IgG1 had the lowest activity against serotype 4 pneumococci [44]. We also found that PMNs were required for production of Abs that protect against infection. This could be mediated by IgG2 [44] or IgG3 which was shown to be protective in mice against pneumococcal infection [45].

How PMNs are inducing class-switching to IgG2c and IgG3 but not IgG1in response to PCV is unclear. Efficient class switching from IgM to IgG requires T cell help [39]. In adults, PCV significantly boosts class switching to IgG as compared to the unconjugated polysaccharide vaccine [12]. Cytokines produced by T cells further determine the subtype of IgG produced with IL-17 and IFN-γ enhancing switching to IgG3 and IgG2 more than IgG1[46, 47]. As PMNs can both produce cytokines [16] and drive Th1 and Th17 cell differentiation [20], they may contribute to isotype switching by producing IFN-γ or IL-17 themselves or eliciting T cells to do so.

In summary, we demonstrate here that PMNs are required at the time of immunization with the pneumococcal conjugate vaccine for optimal protective Ab responses and host protection against subsequent *S. pneumoniae* infection. As serotype replacement by bacterial strains not covered by the current vaccines continue to emerge, novel serotype-independent vaccine formulations such as whole cell vaccines or common pneumococcal protein vaccines are being considered [48]. Therefore, future vaccine designs should take PMN responses into consideration, particularly in susceptible populations like the elderly [2], where PMN responses are known to be dysregulated [49].

## Author Contributions

EYIT conducted research, analyzed data and wrote paper. EAW provided essential reagents and expertise. ENBG designed research, wrote paper and had responsibility for final content. All authors read and approved the final manuscript.

## Acknowledgements

We would like to acknowledge Manmeet Bhalla for critical reading and discussion of the manuscript.

