## Supplemental Materials for "Neutrophils are required during immunization with the pneumococcal conjugate vaccine for protective antibody responses and host defense against infection"

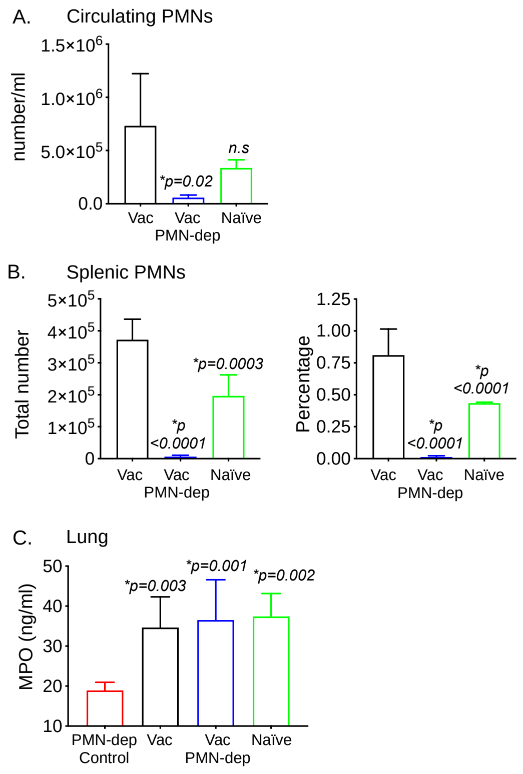


**Supplemental Figure 1. Enumeration of PMNs following depletion and PCV administration.** (A-B) The blood and spleen were collected from naïve, Prevnar-13 immunized and PMN depleted immunized C57BL/6 female mice one week post vaccination following the timeline indicated in Fig 1A. The presence of PMNs (Ly6G+ CD11b+) in the indicated organs was then assessed by flow cytometry. Pooled data from two separate experiments with n=3 naive, n=7 vaccinated isotype treated and n=7 vaccinated PMN depleted mice per group are shown. Bar graphs represent the means+/-SD. Asterisks indicate significant differences with respect to the vaccinated group as calculated by One-way ANOVA followed by Dunnet's test. (C) C57BL/6 female mice were mock-treated (naïve) or vaccinated with Prevnar-13. Vaccinated mice were treated with isotype control or PMN depleting antibodies at the time of vaccination as indicated in Fig 1A. Four weeks following vaccination, the mice were challenged i.t. with $1x10^7$ CFU *S. pneumoniae* TIGR4. Twenty-four hours following pulmonary infection MPO levels in lung homogenates were measured by ELISA. To confirm that PMNs were the source of MPO, a group of vaccinated mice were treated with PMN depleting antibodies one day prior to i.t. infection (PMN-dep control). Data are pooled from two separate experiments with n=6 mice per group. Bars represent the means+/-SD and asterisks indicate significant differences with respect to the PMN-dep control group as calculated by One-way ANOVA followed by Dunnet's test.

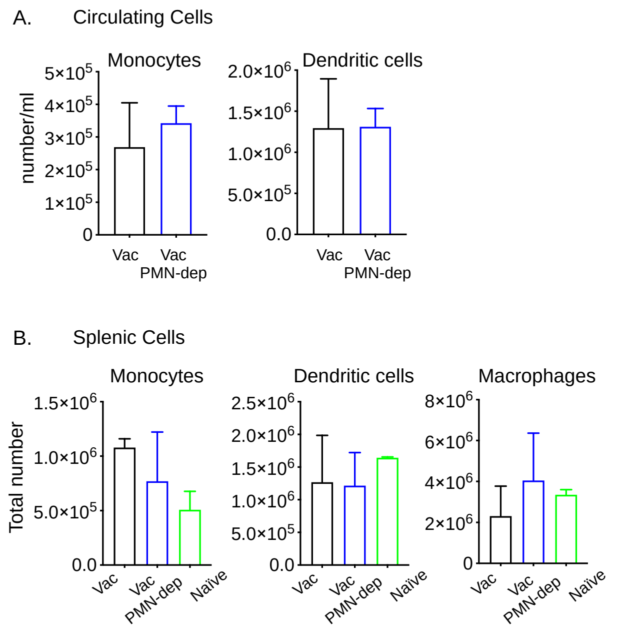


**Supplemental Figure 2. Characterizing circulating and splenic immune cell subsets following PMN depletion.** (A-B) The blood and spleen were collected from naïve, Prevnar-13 immunized and PMN depleted immunized C57BL/6 female mice one week post vaccination following the timeline indicated in Fig 1A. The presence of monocytes (Ly6G$^{-}$, Ly6C$^{hi}$, CD11b$^{+}$), macrophages (Ly6G$^{-}$, F480$^{+}$) and dendritic cells (Ly6G$^{-}$, F480$^{-}$, CD11c$^{+}$) in the indicated organs was then assessed by flow cytometry. Pooled data from two separate experiments with n=3 naive, n=7 vaccinated isotype treated and n=7 vaccinated PMN depleted mice per group are shown. Bar graphs represent the means+/-SD.

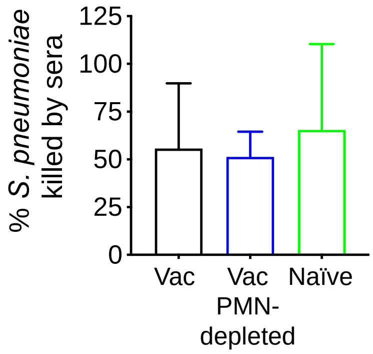


**Supplemental Figure 3. PMN depletion has no direct effect on the anti-bacterial activity of the sera.** *S. pneumoniae* were incubated with sera only from vaccinated, vaccinated PMN depleted or naïve mice for 40 minutes at 37°C. Viable bacteria were enumerated by plating on blood agar plates. The percent of bacteria killed was calculated with respect to a no sera control. Data shown represent the means +/- SD and are pooled from four separate experiments.
